# A non-toxic, user-friendly buffer that enhances green fluorophore performance in conventional and super-resolution imaging

**DOI:** 10.1101/2025.03.27.645765

**Authors:** Anaïs C. Bourges, Mathias Marcos de Vargas, Azevedo, Viktorija Glembockyte, Robin Van den Eynde, Wim Vandenberg

**Affiliations:** Department of Chemistry, KU Leuven, Belgium; Max Planck Institute for Medical Research, Germany

## Abstract

Fluorophore brightness and photostability continue to be limiting factors in fluorescence microscopy. Several buffers that enhance these properties have been identified to date, though they do not work for all dyes, may require harmful components, or have limited storage lives. We have identified a simple, inexpensive, and non-toxic buffer that strongly enhances the brightness and resistance to photodestruction of green emissing synthetic fluorophores, using potassium iodide as a main photostabilizing additive and MgCl_2_ as enhancer for binding kinetics in DNA-PAINT super resolution imaging. We show that this ‘ANice’ buffer can deliver an up to 30% improvement in fluorescence brightness and 5x more detected events in DNA-PAINT imaging. Furthermore, the same effect can be used to strongly increase the photobleaching resistance of all the green dyes tested for conventional imaging.

## Introduction

Fluorescence imaging has become a ubiquitous tool in the materials and life sciences, enabling highly sensitive and specific imaging for a broad range of systems. Most of its applications require labeling the sample with fluorophores, which should typically display a high brightness and photostability so that a strong signal can be observed. Fluorophore brightness and stability are concerns in nearly all applications of this technique, but they are especially pertinent in techniques that rely on single-molecule imaging, such as single-molecule localization microscopy (SMLM). Since brightness and stability are directly related to the achievable detection efficiency as well as the precision of the result.

Over the years, intensive efforts have been dedicated to developing brighter and/or more photostable fluorophores [1]. It was also discovered that the performance of many dyes can be enhanced by the addition of chemical additives, leading to the creation of various imaging buffers tailored to specific fluorophores or applications [2–9]. Such buffers can lead to remarkable gains, promoting blinking behavior of the dyes essential for super-resolution microscopy or reducing photobleaching or transient dark states to increase the number of detectable photons. However, many of these contain toxic components and may have a limited shelf life, rendering them challenging to use. Furthermore, the available imaging buffers are frequently optimized for (far-) red or orange emitting fluorophores, but show less or no enhancement for fluorophores occupying the green region of the spectrum [8]. This renders commonly used fluorophores unsuitable for demanding imaging applications and reduces the possibilities for multicolor or, more generally, high-content imaging in such experiments.

In this contribution, we develop a non-toxic and inexpensive imaging buffer that enhances green-emitting fluorophores photostability and/or brightness. We demonstrate how this ‘ANice’ buffer results in improved SMLM measurements using DNA-PAINT and permits a higher photostability in measurements on permeabilized samples or live-cell measurements in which the fluorophore is displayed on the outer membrane of the cell.

## Results and Discussion

### A new buffer system for increased DNA-PAINT performance of green fluorophores

We previously developed a system simplifying simultaneous multicolor imaging based on point-spread function engineering [10]. While testing the SMLM performance using DNA-PAINT with green-, orange-, and red-emitting fluorophores, we found that the green emitter systematically showed much lower performance than the other fluorophores. A fact that is widely recognized in the field and has been independently verified for a broad range of green-emitting fluorophores [11]. While strategies exist for circumventing this problem by multiplexing red or orange labels using subtle differences in their spectroscopic properties [12–15], achieving a good classification using these methods can be considerably more challenging. This led us to evaluate a set of buffer additives to enhance the performance of the green emitters, prioritizing compounds with low cost, low toxicity, and high chemical stability. In the end, we settled on magnesium and iodide ions. Where Magnesium (in the form of MgCl_2_) is known to function as a kinetic tuner, influencing binding kinetics in DNA-PAINT[16] whereas iodide (in the form of potassium iodide, KI) has been shown to enhance performance of blue-green dyes in fluorescence correlation spectroscopy due to depopulation of triplet and/or radical states [17].

We initially focused our efforts on the green dye Abberior STAR 488 (Ab488), which we had previously selected as the best-performing dye for our SMLM imaging [10]. Here we performed DNA-PAINT experiments with varying amounts of MgCl_2_ and KI added to the commonly used buffer C [18]. The performance was characterized using two parameters: the number of detected blinking events over a defined number of frames and the average detected intensity of these events. In figure 1A, we see that adding iodide increases the apparent brightness up to concentrations of 8 mM while causing a distinct drop beyond this point. The number of detected events also strongly increased when adding iodide, reaching a maximum of 32 mM. The addition of MgCl_2_ gave a slight but noticeable increase in both parameters at increasing concentrations, as seen in Figure 1A. Due to their complementary mechanism, a combination of both compounds was evaluated, where similar trends were seen (data not shown). This led us to propose a combination of 300 mM MgCl_2_ and 8 mM KI in Buffer C as the optimal combination. The straightforward preparation of this buffer, combined with the fact that it only uses comparatively cheap and non-toxic components, led us to refer to this specific composition as buffer “ANice”. When compared to buffer C, buffer ANice leads to a 35% increase in fluorescence intensity and a more than 5-fold increase in detected localization events, even outperforming the PPT buffer system [19], commonly used to enhance dye performance in DNA-PAINT [18] (Figure. 1B and Supporting Movie S1). The increased performance metrics of this buffer also directly translates in the increased quality of the SMLM images, including increased continuity of the stained fibers as well as a reduction of the average localization error (Figure 1C and reconstructed images). Further attempts to increase the performance of the buffer by combining it with oxygen scavengers and triplet quenchers led to no noticeable improvement as shown in the Supporting Figure S1.

**Figure 1:**
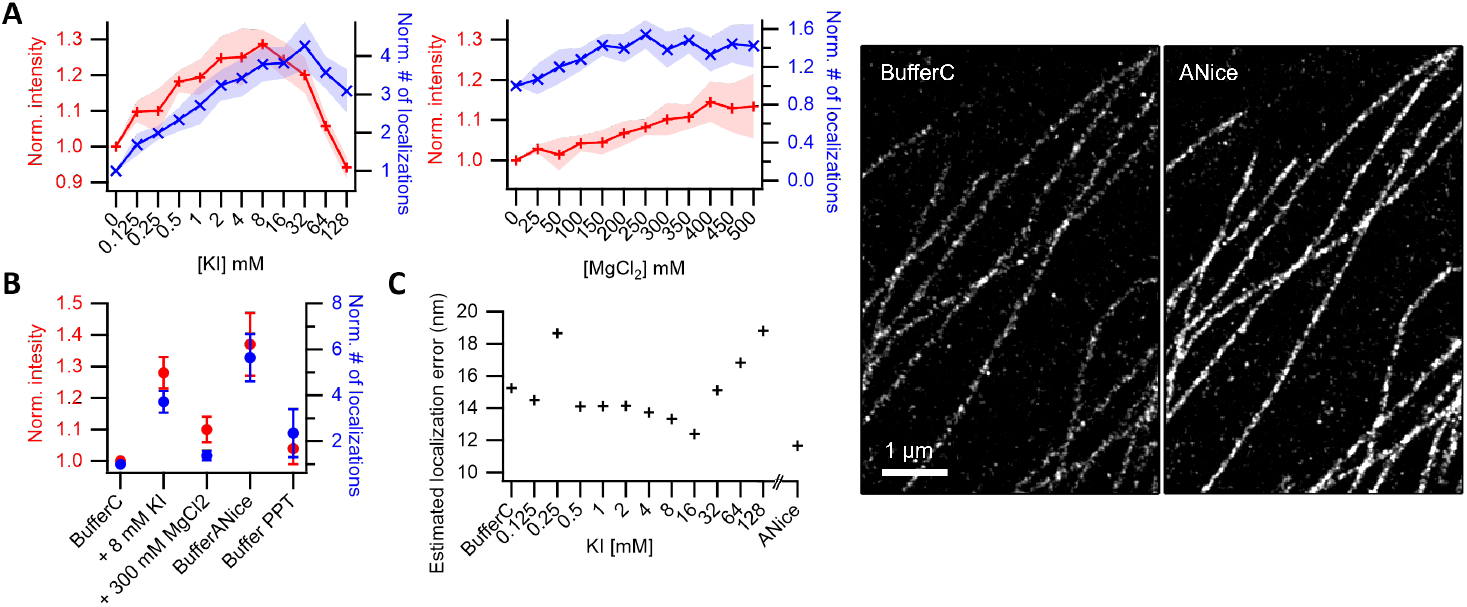
The effect of KI and MgCl_2_ on the DNA-PAINT performance of Abberior STAR 488 (Ab488). A) The amount of detected events (blue) and their average brightness (red) compared to buffer C for varying concentrations of KI and MgCl_2_ (left and right panels, respectively). B) Performance comparison (Number of molecules and Brightness) between Buffer C, Buffer ANice, and PPT. C) Plot the associated localization errors of a KI titration. Right side: Reconstructed image of DNA-PAINT measurements in Buffer C and Buffer ANice.

### Compatibility with orange and red fluorophores

Since one of our goals is to leverage the improved performance of green fluorophores to facilitate simultaneous 3-color DNA-PAINT experiments, we subsequently evaluated the effect of buffer ANice on the Cy3b and ATTO643 dyes that are spectrally compatible with Ab488 [10]. We also assessed the performance of ATTO488, which can serve as an alternative for Ab488. In Figure 2 it can be seen that, at concentrations present in buffer ANice, ATTO488 has a performance increase similar to that of Ab488 while the performance for the red shifted dyes is only slightly altered, with the adverse effects expected from the fluorescence quenching behavior of iodide only occurring at higher concentrations (Supporting Figure S2). In any case, even given slight decreases in performance in Buffer ANice, the localization precision achievable with the orange and red fluorophores still much surpassed the results of Ab488. In contrast, the increased performance of the latter in this new buffer makes 3-color experiments with a sub-10 nm resolution possible (Supporting Figure 3).

**Figure 2:**
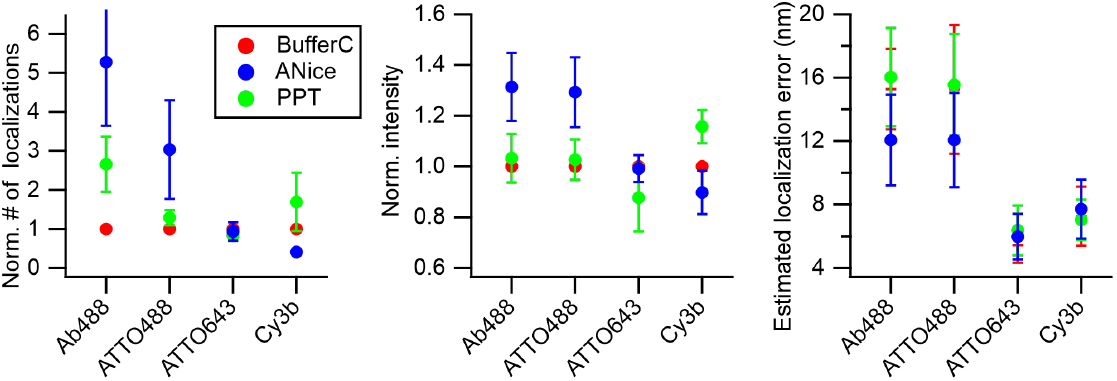
Comparison of the buffer C, PPT and buffer ANice for Ab488, ATTO488, Cy3b and ATTO643 normalized to the performance in Buffer C. For each dye, the number of all the particles detected (left panel), their average intensity (middle panel), and the estimated localization error of the reconstructed image (right panel) are shown.

**Figure 3:**
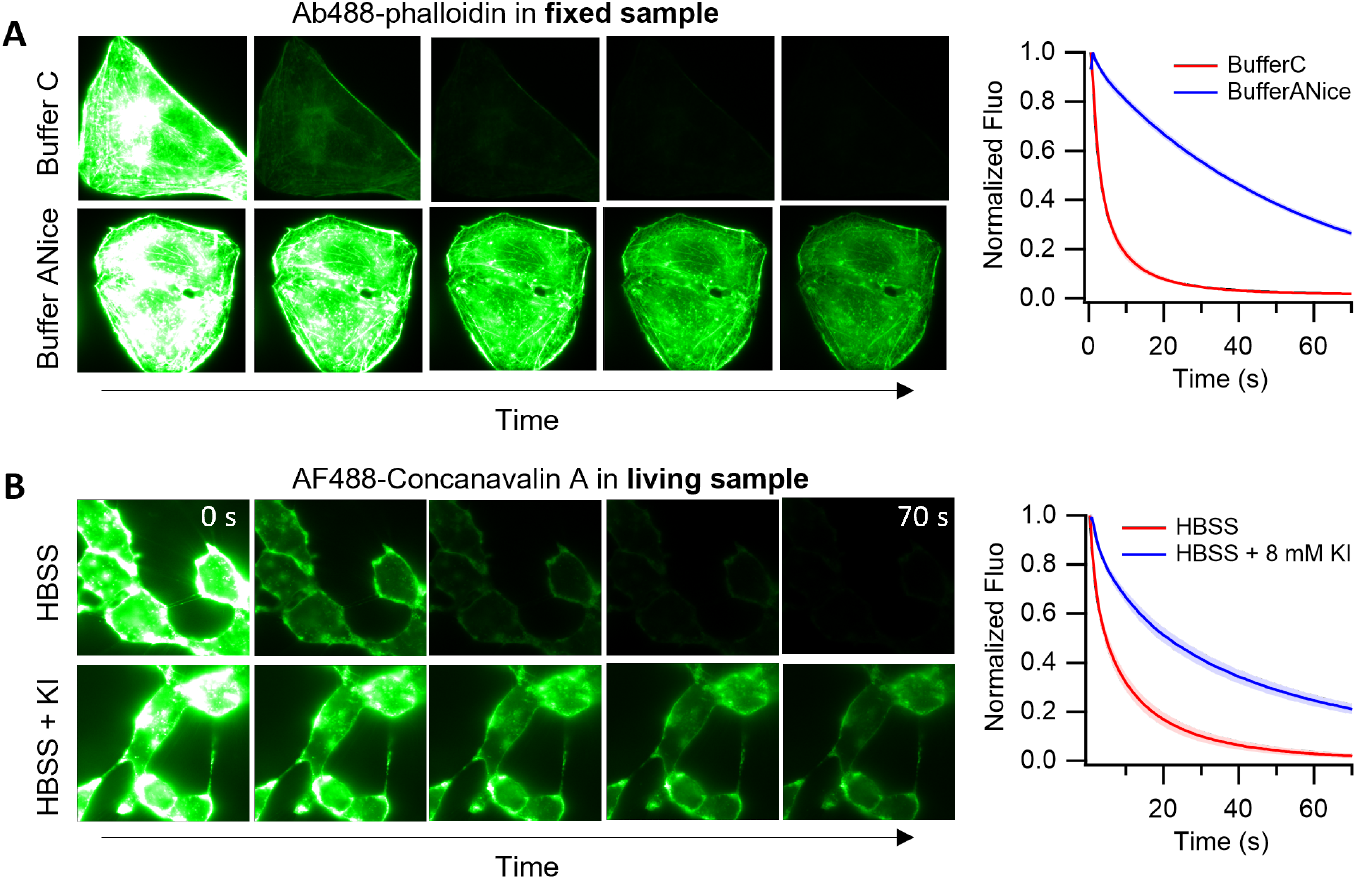
The effect of buffer composition on fluorescence photobleaching. A field of view (FOV) was irradiated several times, with images taken at a regular interval. representative images are shown as well as the average fluorescence intensity traces of 8 FOVs normalized to the initial image. Standard deviations correspond to the light shaded regions. A) Fixed U2OS Cell stained with Ab488-phalloidin imaged in buffer C or buffer ANice. B) Living U2OS cells stained with concanavalin A-AF488imaged in HBSS or HBSS supplemented with 8mM of KI (blue).

### Mechanism of KI on Ab488

We hypothesize that the positive effects of KI stem from either its ability to depopulate the triplet excited state by functioning as a collisional quencher, its ability to recover the dye from a photooxidized dark radical cation state via its redox properties, or a combination of both, as seen in previous work [17]. To identify which of these mechanisms is present in our system, we chose to examine the effects of the reducing agents ascorbic acid (AA) and n-propyl gallate (nPG), on the DNA-PAINT performance (Supporting Figure S4). At micromolar concentrations, these chemicals have been shown to depopulate the majority of photo-oxidized radicals [2, 3], so any additional effect observable when combining them with KI (higher brightness and number of localization) should come from the ability of the later to depopulate the triplet excited states (Supporting Figure S4). Furthermore, any such additive effect should disappear at higher concentrations of AA and nPG, since the reducing agents themselves will depopulate the triplet state at this point. In our experiment, we indeed found this exact signature. Additionally, since a combination of KI and the reducing agents did not additionally increase the performance of the system over KI alone, we believe that in this case KI is once again functioning through both mechanisms. We attribute the effectiveness of the triplet quenching effect to the difference in lifetime of the triplet and singlet excited states in these green emissive fluorophores, since at a concentration of 8 mM singlet excited state quenching is negligible (Supplementary Figure S4) [17]. It is worthwhile to consider that, for the class of green-emissive dyes studied here, the addition of reducing agents alone is sufficient to enhance the optical performance in contrast to the ROXS scheme, which is used for orange- and red-emissive dyes requiring both reducing and oxidizing agents[20].

### Improved resistance to photobleaching in conventional imaging

Spurred on by our results and similar reported improvements [17], we further evaluated whether the enhancement we saw could also benefit conventional microscopy applications. Therefore, we conducted photobleaching experiments on fixed samples, testing a variety of dyes. Here, several fields of view of each sample were irradiated in the presence of buffer C, which was then switched with buffer ANice to repeat the measurements with each sample serving as its control. For all the green dyes tested, an increase in the photobleaching resistance is observed with buffer ANice, whereas for all yellow and red emitting dyes, deleterious effects of buffer ANice are observed (Figure 3, Supporting Figure S5 and Movie S1). When examining the concentration dependency of this effect we found that while the resistance to photobleaching, as measured by bleaching half-time, goes up with increasing concentratin of KI the initial fluorescence shows a maximum at 2 mM of KI (Supporting Figure S6), consistent with our results in DNA-PAINT (Figure 1A). Next, we also evaluated if similar results could be obtained in living cells; for this purpose, the conventional imaging medium (Hanks Balanced Salt Solution, HBSS) was supplemented with 8 mM KI. The effect on cell viability of this concentration was minimal since we found that cells could be readily cultured over several days in media containing this amount of iodide (data not shown). Indeed, for membrane staining using Concanavalin A AF488, a significant decrease in photobleaching rate can be observed (Figure 3B). A similar effect was not seen for intracellular stains, which indicates that iodide is not membrane permeable (data not shown).

## Conclusion

We have shown that adding potassium iodide and magnesium chloride to the imaging buffer significantly improves the DNA-PAINT performance (both number of localization events and their brightness) of green dyes through a synergistic effect of the two compounds. Our improved buffer results in a more comparable performance over the different color channels in three-color super-resolution imaging. Our buffer is easy to prepare and performs similarly even after being stored at room temperature for months. The same underlying effect can also be used to increase the resistance to photobleaching of green dyes in both fixed samples and living cells if the fluorophore is exposed to the imaging medium, enabling longer measurements.

## Materials and methods

U2OS cells were seeded in 8 well plate (Glass bottom, 80807 Ibidi), 15k cells per well. After 24 h cells were fixed as described in [21].

DNA paint measurements. Labeling of the microtubule: anti-Tubulin alpha-Tubulin clone YL1/2 (Thermo Fisher Scientific) used in a dilution 1 in 200, secondary antibody: custom conjugated polyclonal anti-rat IgG (P1 docking strand) [18]. The DNA imaging strands were coupled to either Aberrior STAR 488, ATTO 488, Cy3b and ATTO 646 by IBA. Data were obtained by default in 250 µL of bufferC (1x PBS pH 7.2 with 500 mM NaCl) with 1.2 nM of Ab488, 200 pM of ATTO488, 40 to 60 pM of Cy3b and 40 to 60 pM of ATTO 646. Fresh 1 M potassium iodide (373651000, thermo scientific) and magnesium chloride (25108.295, AnalaR NormaPUR) solutions were prepared in Buffer C, then diluted and stored at the concentration needed. PPT buffer was prepared as described in [18]. Brieflly the buffer is prepared by mixing stock solutions of protocatechuate 3,4-dioxygenase (PCD), protocatechuic acid (PCA) and trolox shortly before the experiement to final concentrations of 0.15 u/mL, 2.5 mM and 50 mM respectively. Imaging was performed using a TIRF illumination and 100 ms exposure time. All measurements were performed with about 2500 images except for reconstructions were more images were needed. Data were analyzed with in-house code using Igor Pro 8 (WaveMetrics Inc.).

Photobleaching measurements on fixed cells. A blocking steps was performed with PBS supplemented with 3% BSA and 0.2% Triton X-100 for 90 min at RT [21]. Dyes were diluted in PBS supplemented with 3% BSA and the staining of the different cells structure was performed following protocols provided by suppliers: Abberior STAR 488 - phalloidin (ST488-0100-20UG) and Abberior Red - phalloidin form Abberior; ATTO 488 phalloidin (AD488-81), ATTO 643 phalloidin (AD643-81), ATTO 6555 phalloidin (AD655-81) from ATTO-TEC; CF 488A phalloidin (00042-T), Wheat Germ Aggglutinin CF 555 (29076) and Wheat Germ Agglutinin CF 568 (29077) from BIOTIUM; Alexa Fluor 488 phalloidin (A12379) from invitrogen. Actin labeling was performed using - Actin antibody (ab6276, Abcam) and NHS-AF555 (Jena Bioscience) according to[22].

Photobleaching measurements on live cells. u2os cells were seeded in 35 mm dish (Glass bottom N1.5, P35G-1.5-14-C, MATTEK) Concanavalin A, Alexa Fluor 488 conjugate (C11252, invitrogen) was diluted in the HBSS buffer to a final concentration of 2 µg mL^−1^ and added to the sample for 30 min at 37C. Wheat Germ Aggglutinin, CF 555 and CF 568 (29076 and 29077, Biotum) were diluted in HBSS to a final concentration of 1 μg/ml and added to the sample for 10 min at RT.

Widefield imaging was performed on a Nikon Eclipse Ti-2 Inverted Microscope (Minato City, Japan) with 1.4 NA oil immersion objective (100× CFI Apochromat Total Internal Reflection Fluorescence) and a ZT405/488/561/640rpcv2 dichroic mirror (Chroma Technology, Vermont, USA). Lasers at 488, 561 and 648 nm (Oxxius, Lannion, France) were used depending on the dyes, as well as the emission filters ET525/50nm (Chroma Technology), Semrock FF01-593/46 and FF01-698/70 (IDEX Health and Science, NY, USA). Images were acquired on a PCO.edge 4.2 camera (PCO, Kelheim, Germany). DNA paint experiments were performed in TIRF illumination with an exposure time of 100 ms and a pixel size of 118.18 nm or 78.8 nm for the estimation of the localization precision.

Photobleaching experiments were performed in epi illumination with an exposure time of 20 ms and a pixel size of 118.18 nm. An initial image was taking followed by an irradiation step of the sample for 0.5 s with the required laser. The laser power was chosen depending on the dye used and its photobleaching rate. These steps were repeated multiples times for different FOVs of the sample. If necessary, the sample was washed with PBS before the next buffer was added. One or multiple cells were selected per FOVs for analyses and a background subtraction step was done before normalizing the fluorescence intensity curves to the initial (first) image of the stack.

Absorbance and fluorescence spectra were measured on a homemade system[23] in a cuvette. The dye Ab488 was diluted in 100 µL ul to a final concentration of 2 μM in buffer C. For each concentration of KI, the desired amount of KI solution at 1 M was added. Later a dilution factor was applied to the analyzed spectra.

## Acknowledgments

A.C.B. thanks the European Commission for a Marie Skłodowska-Curie postdoctoral fellowship. This work was supported through funding from the Research Foundation Flanders through grants 1514319N, G090819N, G0B8817N, the KU Leuven through grant STG/22/059, and the European Research Council through grant 714688 NanoCellActivity. Peter Dedecker is thanked for valuable input and discussion.

