## Supporting information for "A non-toxic, user-friendly buffer that enhances green fluorophore performance in conventional and super-resolution imaging"

#### 1 Supporting movies

Supporting Movie S1: Comparison of the performance of the Abberior STAR 488 (Ab488) fluorophore in buffer C and buffer ANice. Upper panels: DNA-PAINT experiment of the same region with exactly the same parameters, except that buffer C was replaced with the buffer ANice for the video on the right. Lower panels: Photobleaching experiments of fixed cells stained with phalloidin-Ab488. The same parameters were used except for the buffer.

#### 2 Comparison to existing buffer systems

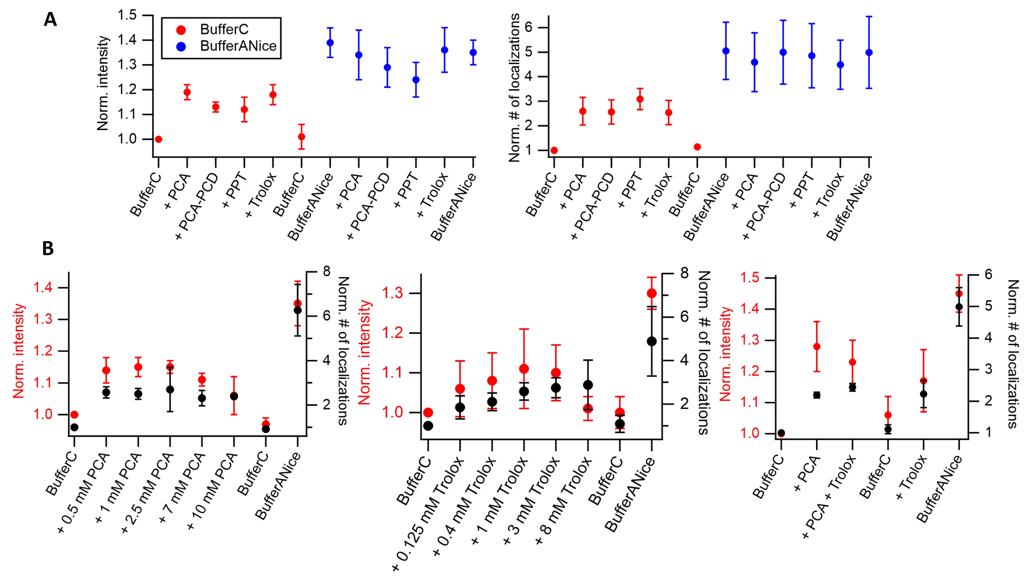


Supporting Figure S 1: The DNA-PAINT performance of Ab488 for combinations of the PPT and ANice buffer system normalized to the performance in Buffer C, quantified by average spot brightness and number of detected localizations. A) the effect of individual PPT components alone or in combination when added to Buffer C or buffer ANice. B) The effect of different concentrations of PCA and Trolox in buffer C. For each independent measurement of a FOV, the buffer was replaced by the following one in the same order as presented in each pannel.

#### 3 Performance of 3 color DNA-PAINT in function of iodide and magnesium ion concentration

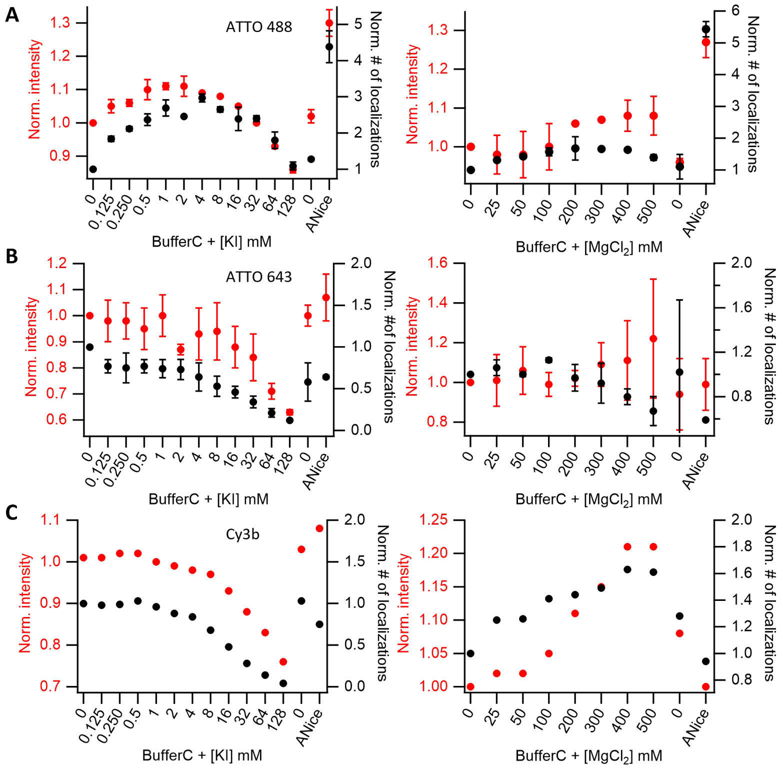


Supporting Figure S 2: The DNA-PAINT performance of ATTO488, Cy3b and ATTO643 for varying concentrations of KI and $MgCl_2$ normalized to the performance in Buffer C quantified by average spot brightness and number of detected localizations.

### 4 Performance of 3 color DNA-PAINT in buffer ANice

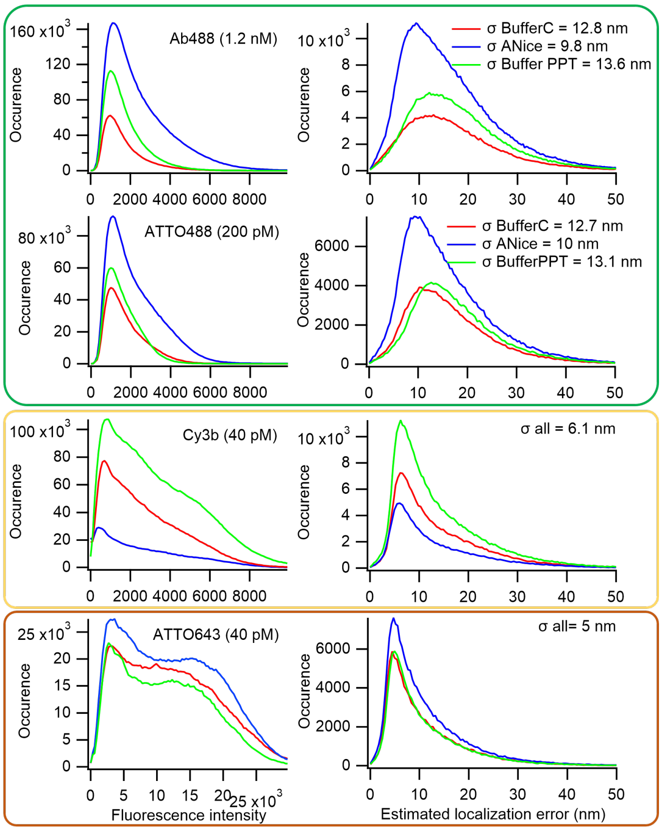


Supporting Figure S 3: Histogram of detected brightness and estimated localization precision for DNA-PAINT experiments of Abberior STAR 488 (Ab488), ATTO488, Cy3b and ATTO643 in Buffer C, PPT and Buffer ANice. Dyes with similar emission color are grouped in the same frame.

### 5 Proposed mechanism of KI

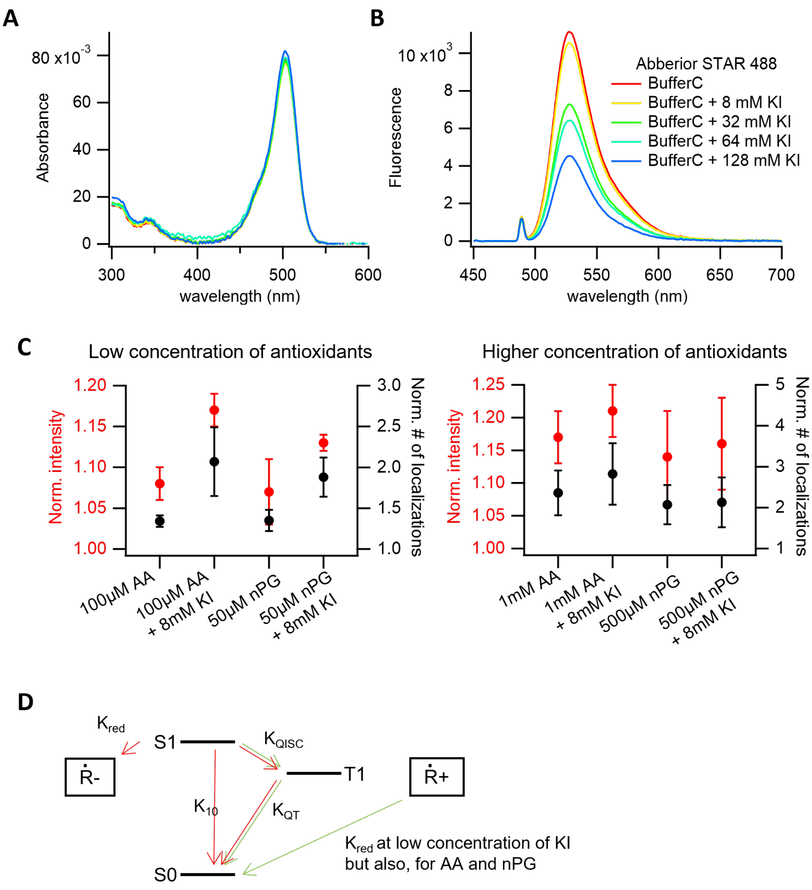


Supporting Figure S 4: Mechanism for KI. A) Absorbance and B) emission fluorescence of Abberior STAR 488 (Ab488) in buffer C supplemented with increasing concentrations of KI measured in a cuvette. C) DNA-PAINT measurements of Ab488 in the presence of ascorbic acid (AA) or n-propyl gallate (nPG) and potassium iodide (KI) in buffer C. At low concentrations of AA or nPG (left panel), the increases in intensity and the number of detected localizations could come from KI depopulating the triplet states as most of the free radical are reduced by the 2 antioxidants AA and nPG. At a higher concentration of AA and nPG, no additional effects are seen. Values were always normalized to the first data point of a set of measurements on the same FOV, corresponding to buffer C. D) A simple representation of the rate constants affected by KI. In green, the positive effect of KI at low concentrations. Note that $K_{red}$ is also positively affected by µM of AA and nPG. Higher concentrations of KI lead to an increase of rate constants inducing an accumulation of dark state species (in red) and a decrease in overall fluorescence as seen in B. $K_{10}$ is the relaxation rate form S0 to S1, $K_{QT}$ correspond to the triple state quenching and $K_{QISC}$ the intersystem crossing, all influenced by KI.

### 6 Photobleaching curves for selected dyes

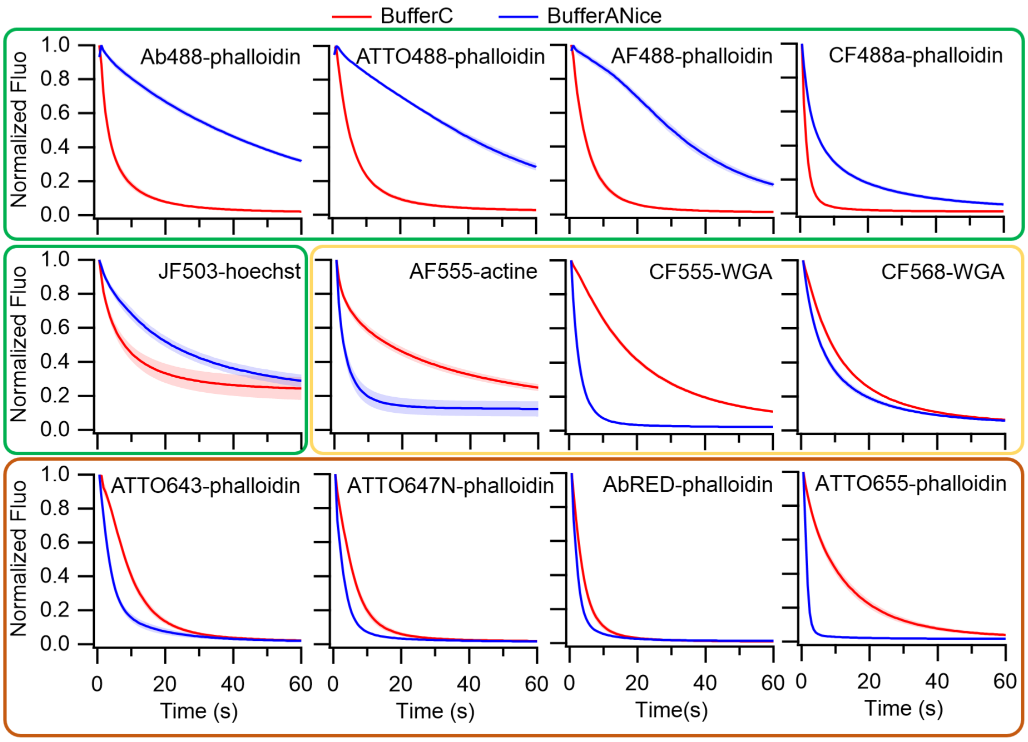


Supporting Figure S 5: Photobleaching of U2OS cells labeled with a variety of dyes. Average fluorescence intensity traces of 8 fields of view normalized to the initial image acquired in buffer C (red) or buffer ANice (blue). Standard deviations correspond to light shaded regions. Dyes with similar emission color are grouped in the same frame.

### 7 The effect of iodide ion concentration of photobleaching of Aberior Star 488

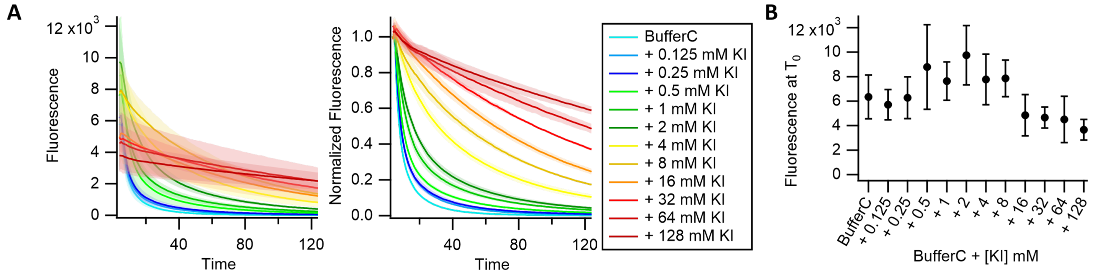


Supporting Figure S 6: KI titration of fixed U2OS cells labeled with Ab488-phalloidin. A) A field of view (FOV) was irradiated several time for 0.5 s each time followed by an image with an 20 ms acquisition time. Average fluorescence intensity traces of 5 FOVs (left panel) and normalized to the initial image (before any irradiation step, right panel) acquired in Buffer C or buffer C supplemented with increasing concentrations of KI. Standard deviations correspond to light shaded regions. B) Average fluorescence of the first image from the data show in A.
